# Reticulate evolution and rapid development of reproductive barriers upon secondary contact pose challenges for species delineation in a forest fungus

**DOI:** 10.1101/2023.09.18.558330

**Authors:** Dabao Sun Lu, David Peris, Jørn Henrik Sønstebø, Timothy Y. James, Loren H. Rieseberg, Sundy Maurice, Håvard Kauserud, Mark Ravinet, Inger Skrede

**Affiliations:** Department of Biosciences, University of Oslo, Oslo, Norway; Department of Biotechnology, Institute of Agrochemistry and Food Biotechnology (IATA), CSIS, Paterna (Valencia), Spain; Department of Natural Sciences and Environmental Health, University of South-Eastern Norway, Bø, Norway; Department of Ecology and Evolutionary Biology, University of Michigan, Ann Arbor, USA; Department of Botany and Biodiversity Research Centre, The University of British Columbia, Vancouver, Canada; School of Life Sciences, University of Nottingham, Nottingham, UK

## Abstract

Reproductive barriers within morphospecies of the mushroom-forming fungi tend to be stronger between sympatric lineages, leading to speculation on whether they are being reinforced by selection against hybrids. In this study we use population genomic analyses together with *in vitro* crosses of a global sample of the wood decay fungus *Trichaptum abietinum* to investigate reproductive barriers within this morphotaxon and the processes that have shaped them. Our phylogeographic analyses show that the morphospecies is delimited into six major lineages, one in Asia, two in Europe, and three in North America. The two lineages present in Europe are interfertile and admixed, whereas our crosses show that the North American lineages are reproductively isolated. In Asia a more complex pattern appears, with partial intersterility between multiple sub-lineages that likely originated independently and more recently than the reproductive barriers in North America. We found pre-mating barriers in *T. abietinum* to be moderately correlated with genomic divergence, whereas fitness of the mated hybrids showed a strong positive correlation with increasing genomic divergence. Genome wide association analyses identified candidate genes with programmed cell death annotations, which are known to be involved in intersterility in distantly related fungi. Our demographic modelling and phylogenetic network analyses suggest that reproductive barriers in *Trichaptum abietinum* could have been reinforced upon secondary contact between lineages that diverged in allopatry during the Pleistocene glacial cycles. Our combination of experimental and genomic approaches demonstrate how *T. abietinum* is a tractable system for studying speciation mechanims.

## Introduction

It is estimated that kingdom fungi contains between 1.5 and 7 million species^1–3^, yet our knowledge about the mechanisms that have generated this diversity is disproportionally small in comparison to what we know about animals and plants. Unique life cycles, small genomes, and a relatively high vagility may set speciation mechanisms in fungi apart from those of other eukaryotes^4–6^. The development of reproductive barriers is central to the speciation process as they restrict geneflow between populations and thereby fortify them as independently evolving lineages^7^. Work on fungal model organisms^8,9^, and pathogens of plants^10^ and humans^11^, have found that reproductive barriers in fungi can be caused by genetic incompatibilities^12,13^, chromosomal rearrangements^14,15^ and ecological specialization^16,17^. Another mechanism that can create strong reproductive isolation is reinforcement. Here natural selection works directly against maladaptive hybridization by favoring pre-mating barriers^18^, and consequently these are expected to be stronger between populations in sympatry than allopatry^19^. In compiling information on reproductive barriers in fungi, Le Gac and Giraud^20^ found pre-mating barriers to be significantly stronger for species pairs in sympatry than allopatry in subclass Agaricomycetes^21^, suggesting a role for reinforcement in shaping the reproductive barriers in this group, which contains the mushroom forming fungi.

Many agaricomycete wood decay fungi comprise species complexes with wide distributions and intersterility groups (ISGs) in sympatry^22–27^. In addition, they are easy to culture and cross in the laboratory, which makes them suitable for investigating fungal speciation and reinforcement. In this study we focus on *Trichaptum abietinum* (Pers. ex J.F. Gmel.) Ryvarden, where previous crossing experiments have revealed two sympatric ISGs in North America, and a third European ISG which can mate with both of the North American ISGs^23,28^. *Trichaptum abietinum* is a common decay fungus on dead wood of tree species of *Pinaceae* across boreal and temperate regions. Molecular studies of rDNA^29–31^ and genome data^32^ have revealed multiple divergent lineages within *T. abietinum*, and two of these have been connected to the ISGs previously detected in North America^31^.

In this study we use population genomic analyses in conjunction with lab experiments on a global collection of *T. abietinum* to learn more about the processes that have generated the high species diversity in fungi. First, we apply a phylogeographic framework to assess how many lineages are present globally. We predict that the genetic structure reflects survival in different refugia during the last glacial maximum, but to a lesser extent than in animals and plants due to a high vagility. We then test and compare pre- and post-mating barriers between lineages using crossing and growth experiments, where stronger pre-mating barriers between sympatric lineages can be interpreted as support for reinforcement, although we recognize that other evolutionary processes can produce a similar pattern^19^. We search for the genetic basis of intersterility between lineages with genome scans and genome wide association studies, and expect barrier loci involved in reinforcement selection to show a signature of positive selection, similar to what has been shown in other systems^33–36^. Secondary contact is generally a precondition for reinforcement^18^, and we ask if the lineages with pre-mating reproductive barriers show evidence of this.

## Results and Discussion

### Phylogeography detects six major lineages within the *Trichaptum abietinun* morphotaxon

To uncover the global diversity and genetic structure within the *T. abietinum* morphotaxon, we obtained fruit body samples covering most of its distribution (suppl.table1). The genomes of 358 haploid (monokaryotic) cultures from these samples were sequenced to an average depth of 24x on an Illumina platform and mapped to a reference genome of *T. abietinum*^32^ to obtain a SNP dataset for population genomic analyses. We used three different methods to assess the population structure: i) Principal component analysis (PCA) ii) maximum likelihood (ML) phylogeny and iii) Admixture analysis, which in addition to population structure can detect admixture, i.e. when samples contain ancestry from multiple lineages. Altogether, these analyses support the division of *T. abietinum* into six major lineages. Our PCA showed five distinct lineages: three in North America, one in Europe and Siberia, and one in East Asia (Fig. 1a). North America C consisted of only two samples from higher elevation sites in Western North America, and is likely underrepresented in our study as these areas were not sampled extensively. The same five lineages were detected in the ML phylogeny (Fig. 1b), whereas Admixture^37^ split the European samples into two lineages; one consisting of samples from southern Europe and another containing samples from Northern and Eastern Europe extending into Siberia, hereafter referred to as the European and Eurasian lineages, respectively (Fig. 1c, 1d). A number of samples from Europe and East Asia appeared admixed, and we will discuss these results below (see sections on reproductive barriers and secondary contact). North America C appeared as an admixture of North America A and the European lineage, which may be an artefact caused by the low sample size^38^. In comparing the ITS sequences of our samples to those of Seierstad et al.^31^, we determined that North America A and B correspond to the previously detected ISGs of Macrae^23^. While Seierstad et al.^31^ inferred a widespread Circumboreal lineage in *T. abietinum* covering both Europe, Asia and North America using rDNA markers, our genome data from a similar spread of populations shows that this group includes three differentiated and geographically restricted lineages; Eurasian, East Asian, and North America B, all with Fst ≥ 0.2 (suppl.table2).

**Figure 1.**
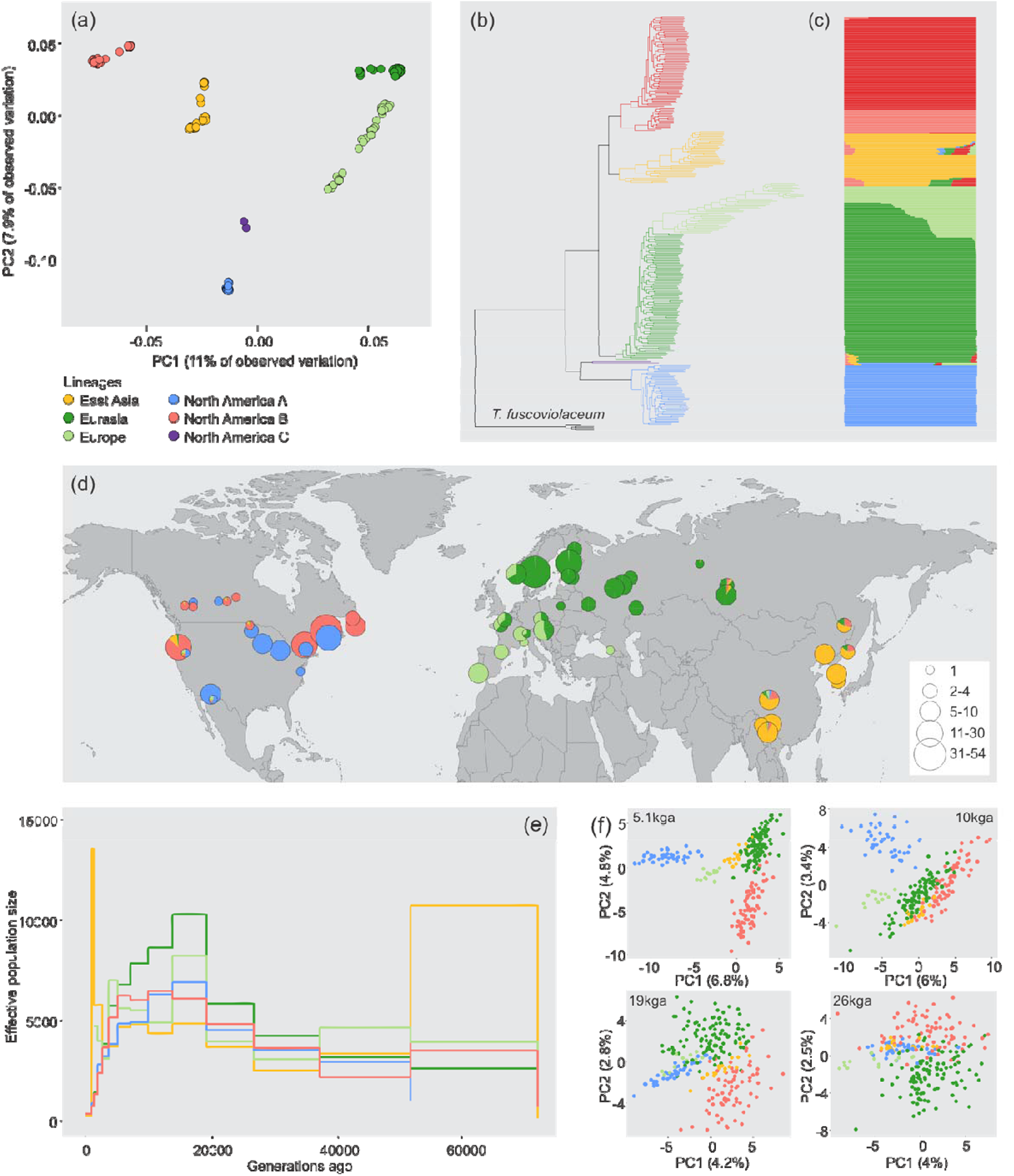
Phylogeographic patterns of Trichaptum abietinum based on population genomics data **a**) Principal component analysis (PCA) showing five distinct clusters. A sixth distinct European lineage detected in other analyses is colored in light green. **b**) Maximum likelihood (ML) phylogeny including the sister species T. fuscoviolaceum as outgroup showing the same 5 major lineages as in the PCA, with 100% support values for all major lineages **c**) Admixture analysis of K = 6, which had the lowest cross validation error. The samples are ordered in the same order as the ML phylogeny, and identify an additional cluster of North America B from western North America. **d**) Populations plotted with their average admixture proportions on a world map. The size of the circle indicates the number of samples from the population. If two lineages are found at the same site there are two circles i.e. North America A and B in eastern North America and North America C with North America A and B in western US. e) Coalescence analysis showing estimated effective population size through time for selected populations of 5 of the lineages (excluding North America C). f) PCA of the pairwise coalescence rate for the same samples as in e) at selected time points. Color denote lineage the sample have been ascribed to.

We performed coalescence analyses in Relate^39^ to infer historic relations between the major lineages and their changes in effective population size (Ne). All lineages showed a strong increase in Ne about 20,000 generations ago (Fig. 1e). Since the generation time of *T. abietinum* is unknown, the number of generations cannot readily be converted to years. *Trichaptum abietinum* is a primary colonizer that rapidly fruits on recently dead wood substrates, and we note that the rise in Ne coincides with the end of the last glacial maximum 20,000 years ago if we assume a generation time of one year. The lineages did not coalesce on the graph (Fig. 1e), possibly indicating limitations of this method for the small genome size of 50 Mb for *T. abietinum*^40^. PCAs of the pairwise coalescence rates between all individual samples back in time show that the Eurasian, East Asian and North America B lineages cluster together at several time periods with similar coalescence rates indicating shared ancestry. We interpret this as high levels of gene flow between these lineages, and it is possible that the rise in Ne 20,000 generations ago is due to increased gene flow between all lineages. At any event, the negative Tajimás D values for Eurasia indicate that recent population expansions have occurred here (suppl.table3). Interpreting this in light of the greater similarity in pairwise coalescence with East Asia and North America B (Fig. 1f), we suggest that the Eurasian lineage has an eastern origin. Similarly, we suggest that North America B originated in Beringia from East Asian ancestors, and migrated to North America over the land bridge, but also spread westward to East Asia and Europe, where it mixed back in with existing lineages (Fig. 1f). Southeastern US was a refugium for boreal vegetation when the Laurentide ice sheet covered most of northern North America^41^. We hypothesize that North America A expanded from this refugium considering how they were more frequent in our samples from eastern and southern parts of the continent and that the nucleotide diversity is higher in the south (suppl.table3). Similarly, North America C may have originated south of the Cordilleran ice sheet in western North America. However, further sampling near the historic refugia is needed to verify these hypotheses. Considering how Asia remained mostly ice free during the ice ages^42^, more continuous geneflow between populations could have kept the lineages here from diverging to the same extent as on the other continents. The biogeographic patterns we observed of higher levels of genetic diversity in the south and a greater number lineages in North America, has been found in a number of other Agaricomycetes^22,43–45^. The abovementioned patterns are also known trends for plants and animals^46^, suggesting that these fungi are structured by similar biogeographic processes and could have followed the same postglacial migration routes, but more fine grained studies are needed here.

### Crossing experiments reveal multiple reproductive barriers

Interfertility in Agaricomycetes has been scored by the ability of two monokaryotic mycelia to form clamp connections upon pairing, as the presence of these structures indicate a dikaryotic mycelium which results from a successful mating between monokaryons^23,47,48^ (Fig. 2a). To assess mating barriers within the *T. abietinum* morphospecies, two to five selected monokaryotic isolates from each of the six major lineages were crossed in all combinations (Fig. 2b). In correspondence with previous findings^23,28^, the majority of our crosses between North America A and B produced no clamp connections. The exceptions were four parings between allopatric North America A and B isolates that resulted in sparse production of clamps. All sympatric inter-lineage pairings were completely intersterile, consistent with reinforcement, where stronger pre-mating barriers are expected in sympatry. Our crosses revealed additional ISGs: North America C was completely intersterile with North America A. Surprisingly, the two samples of North America C were not interfertile with each other, and may represent reproductively isolated populations. Moreover, North America A showed almost complete intersterility with the samples from Yunnan and Sichuan in the East Asian lineage. The isolates from Yunnan were also largely intersterile with all the other major lineages. Due to the complex pattern in East Asia, we also crossed 12 isolates from East Asia in all combinations (Fig. 2c). This revealed that the Yunnan population is intersterile with the population from Heilongjiang and partially intersterile with the Sichuan population. Interestingly, both of these populations appear to be admixed (Fig 1d), alluding to a role for hybridization in causing intersterility here. In any event, these barriers must be recent given the low divergence between the populations, suggesting that pre-mating barriers can evolve rapidly. The Eurasian and European lineage were almost entirely interfertile with each other, whereas the other crosses between lineages showed a patchwork of mixed intersterility lacking clear patterns, indicating that intersterility is governed by several loci, similar to what has been inferred in the wood decay fungus *Heterobasidion*^49^.

**Figure 2.**
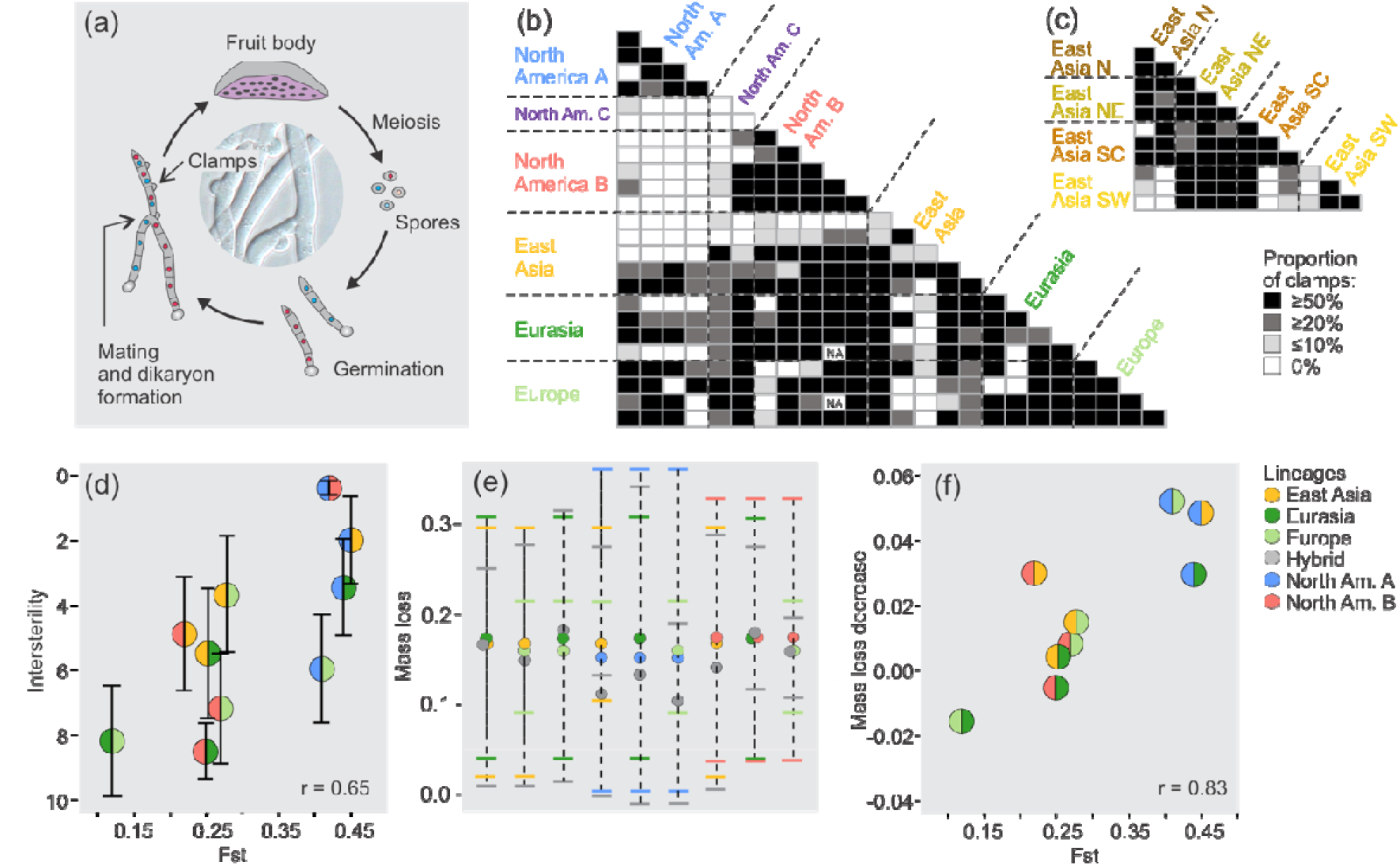
**a**) Overview of typical life cycle for basidiomycetes, including Trichaptum abietinum. Basidiospores produced by meiosis in the fruit body germinate into haploid mycelia known as monokaryons. Interfertile monokaryons can mate and form a dikaryotic mycelium, and this is the dominant stage of the life cycle. Dikaryons can be recognized by clamp structures which extend between cell junctions, and their presence or absence in crosses have been used to score intersterility/interfertility between populations. **b**) Crosses of selected monokaryons from all the major lineages. Crosses were scored according to the average proportion of clamps. For each cross we examined 10 cell junctions for clamps. The majority of crosses were set up in triplicates, thus the % is based on 30 possible clamp occurrences. **c**) Crosses between sublineages in East Asia scored the same way as in b) **d**) Intersterility measured as mean number of clamps for crosses between major lineages plotted against genetic distance between major lineages as measured in average Fst between lineages (r = 0.65, p = 0.04). Correlation calculated using Pearsons correlation test. Bars delimit 95% confidence interval. **e**) Proportion of mass loss of decomposed pine wood blocks by hybrid dikaryons as compared to each parental lineage. **f**) Mean of differences in decomposition in hybrid dikaryons as compared to the average decomposition of each parent lineage plotted against genetic distance between major lineages as measured in average Fst between lineages. (r = 0.83, p = 0.005) Correlation calculated using Pearsons correlation test.

We observed a moderate correlation of 0.65 (p = 0.04) when relating Fst values between the major lineages to their degree of reproductive isolation (Fig. 2d). When including the incompatibility patterns between the 12 East Asian isolates, the correlation dropped to 0.51 (p = 0.04). The general increase of intersterility with genetic divergence we observe fits with speciation in allopatry where reproductive barriers result from genetic incompatibilities, which has been proposed for several Agaricomycetes^27,50,51^. However, sufficient build-up of genetic incompatibilities to cause intersterility may take considerable time^52^, and we note that in contrast to the overall trend, North America A and the Yunnan population in the East Asian lineage were able to mate with isolates that are equally or more divergent than the isolates they cannot mate with in sympatry or parapatry. Similar patterns have been observed in several basidiomycete species complexes^51,53,54^, which suggest these pre-mating barriers are caused by more than the clock-like accumulation of BDMI and may involve reinforcement selection.

Mating tests between monokaryotic cultures only assess prezygotic plasmogamy and do not account for the actual fitness of the dikaryon and its ability to produce a fruit body and complete the life cycle (Fig 2a). As a simple assessment of their fitness, we measured the wood decay capacity of hybrid dikaryons between five of the major lineages, and compared it to that of the dikaryons synthesized from within the lineages. In seven out of nine cases, the hybrid dikaryons decomposed wood slower than dikaryons from parent populations (Fig. 2e, <u>suppl.fig_dikaryon_fitness</u>), and the hybrids between East Asia and North America A showed a significantly lower decay rate than either parental population. Overall, we observed a strong correlation (r = 0.83, p = 0.005) between Fst values of parent populations and the reduction in wood decay of their hybrid dikaryons (Fig. 2f). Our measurements of decay under lab conditions may not reflect how well dikaryons perform in nature, but we note that the correlation is stronger between wood decay ability and Fst than between intersterility and Fst. Lower hybrid fitness has been documented in other basidiomycete fungi^55,56^, and makes a case for reinforcement to avoid investment in unfit hybrids. In the same vein, the increased fitness between hybrids of the Eurasian and European lineage could explain the lack of pre-mating barriers here.

To search for genetic regions associated with intersterility we applied a combination of genome wide association studies and genome scans. We ran a genome wide association study (GWAS) on intersterility with North America A where all samples were included, and intersterility was inferred based on lineage for the samples which we had not included in our crossing experiments. In addition, we conducted a multivariate analysis of correlated phenotypes with SCOPA^57^ where the 25 samples in our crossing experiment and their intersterility with all 6 major lineages were analysed together. Assuming that intersterility in basidiomycetes is caused by heterogenic incompatibility, we searched for genomic regions that are highly differentiated between lineages that cannot mate, but similar in the lineages that can mate, as quantified by ΔFst. Functional annotations of these analyses revealed 407, 1580, and 1357 genes that contained at least one PFAM domain and overlapped with significant GWAS SNPs, SCOPA SNPs, or ΔFst windows, respectively. (Suppl.table4, Suppl.table5, Suppl.table6). The main functions of these lists of genes are difficult to evaluate, and there is a confounding effect of population structure since intersterility is highly correlated with lineage (Suppl.figGWAS). However, there are a few clues on genes to search for; a recent study of the fungus *Podospora* by Ament-Velásquez et al^58^ found genes for somatic incompatibility that regulated cell death to also be causing intersterility between populations. In our dataset, two of the 62 SNPs with 0 p-value in the SCOPA analysis, the 12^th^ most significant GWAS SNP, and 5 of the genes within ΔFst outlier windows were annotated as metacaspases. Metacaspases regulate cell death^59^ and are known to function in somatic incompatibility in the fungal phylum Ascomycota^60^. In total, 7 out of 9 genes putatively annotated as metacaspases turned up in one of these three analyses. The metacaspase on scaffold 6 came up in all three analyses, and was identified as ΔFst outlier among populations of the East Asian lineage, suggesting they may play a role in the partial reproductive barriers here. The genes underlying somatic incompatibility are not characterized in agaricomycetes^61^, but programmed cell death have been predicted to function in somatic compatibility here^62^, and we speculate that this machinery may have been co-opted for intersterility in *T. abietinum*. A test for positive selection in Relate on the increase in frequency of variants in the inferred genealogies did not detect any significant signal for positive selection within the metacaspase gene annotations in North America A, North America B, or the East Asian sublineages, which are the lineages we predict to have experienced reinforcement selection.

### Relating reproductive barriers to secondary contact

We carried out additional analyses to assess if the pre-mating barriers in *T. abietinum* could be linked to secondary contact, and by extension, to reinforcement. To investigate secondary contact, we used Treemix^63^ to construct phylogenies that can account for admixture of lineages. The suggested 99.8 threshold of explained variation^63^ was reached in a phylogenetic network with five admixture events that included admixtures between both the reproductively isolated North America A and B, and the partially reproductively isolated Sichuan and Yunnan populations in East Asia (Fig. 3a). The admixture was estimated as weak between the two North American lineages, suggestive of introgression, but relatively strong in East Asia, consistent with the creation of a new lineage through hybridization. In addition, Treemix identified introgression into the East Asian lineages from an unsampled lineage related to the outgroup *T. fuscoviolaceum*. Kinneberg et al.^64^ found support for ancient introgression between *T. abietinum* and *T. fuscoviolaceum*, and it is conceivable that introgressed genes from the *T. fuscoviolaceum* clade have spread into North America A and B through populations in East Asia.

**Figure 3.**
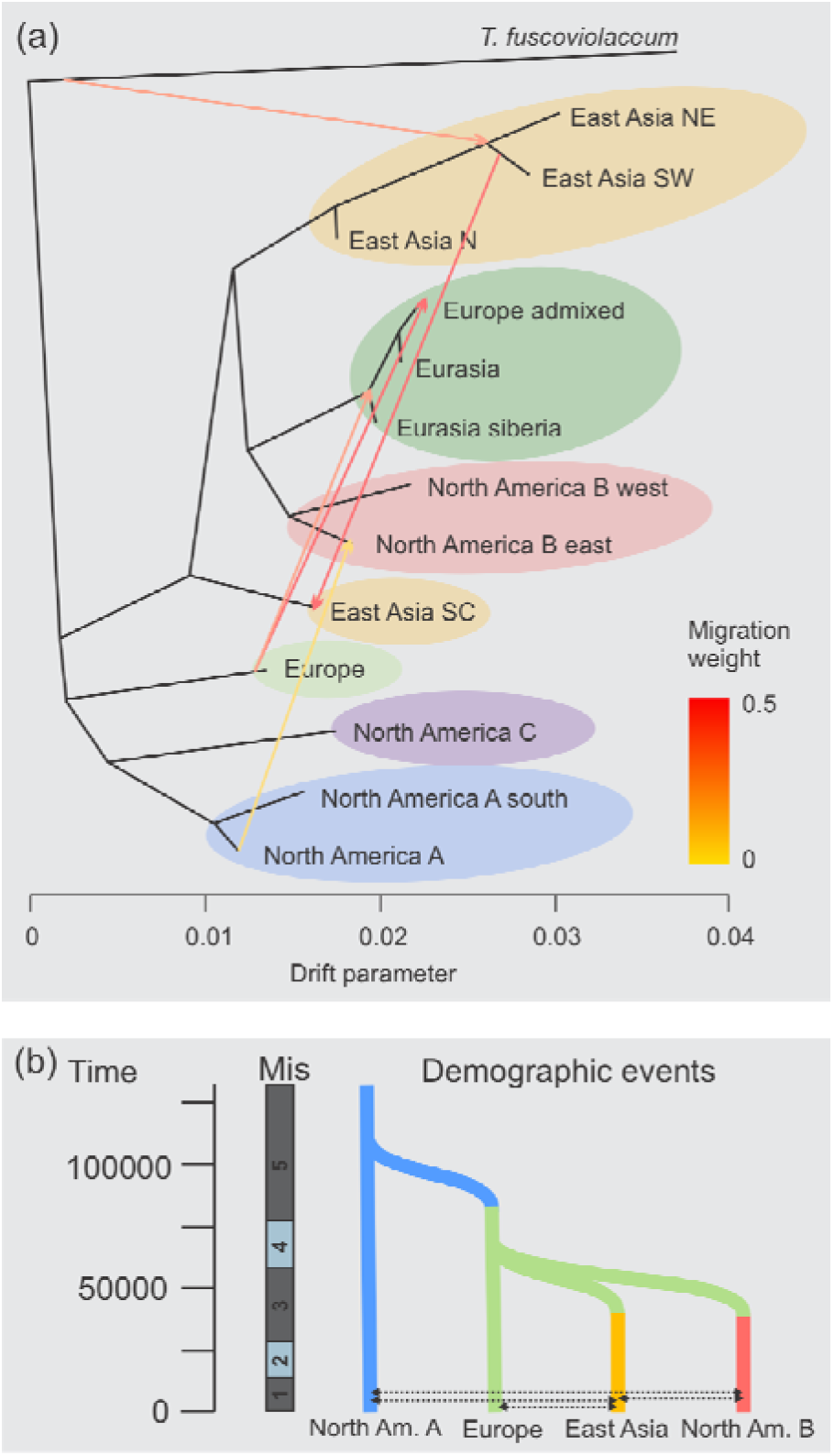
a) Phylogenetic network in Treemix with 5 admixture events, indicated by arrows between lineages and colored by migration weight. b) Coalescence and geneflow events in the best of Fastsimcoal2 model plotted with time, assuming a generation time of 1 year, and major isotopic stages (Mis). Mis captures fluctuations in earth temperature, and cold periods are shown in blue, where Mis2 corresponds to the last glacial maximum.

We applied demographic modelling with Fastsimcoal2^65^ to explicitly test for secondary contact between the reproductively isolated North America A and B, and to estimate divergence time between the major lineages (<u>table1</u>). All models with gene flow were significantly better than the model without gene flow. Among the models with gene flow, those including secondary contact between North America A and B had a significantly better likelihood than comparable models with the same number of parameters (<u>table1</u>, <u>suppl.fig.likelihood_distributions</u>). The best model included four migration events, and here the coalescence time between North America A and the other lineages was estimated to 113,000 generations ago. Coalescence time generally precedes divergence time^40^, and if we assume a generation time of 1 year, the divergence between North America A and the European lineage happened around the end of the previous interglacial^66^. In the same model, North America B and East Asia can be inferred to have diverged from Europe sometime after their coalescence 60,000 generations ago, after the onset of the last ice age but around a minor transition when the global climate experienced some warming (fig. 3b). The best model included geneflow between both East Asia, Europe, and North America A and B, and it is possible that the more recent coalescence of these groups indicated by Relate captured the geneflow that occurred after the main split. It is conceivable that the reproductive barriers between sub-lineages in East Asia and North America A also resulted from reinforcement selection, but they may also be a by-product from geneflow with North America B.

Given the evidence of geneflow between lineages, we conducted a sliding window PCA with Lostruct^67^ to investigate if specific genomic regions stood out with a signal of introgression. The analysis indicated a different pattern from the genome average on the second half of Scaffold 5 (Fig 4a,4b). A PCA plot of this region revealed that the North America A, B, and C lineages to be more similar here when compared to the PCA of the whole genome (Fig. 4c). Inspection of our Fst genome scans showed a similar pattern, where the second half of scaffold 5 had lower divergence between North America A and B than any other Fst comparisons including these lineages (Fig. 4d). This could indicate an inversion that has spread from North America A to B. Inversions are predicted to enhance reinforcement and rapid development of reproductive isolation as they tend to suppress the recombination rate^68^, and we note that two of the metacaspases from the ΔFst outlier windows fall within this region. If the introgression on scaffold 5 was part of the secondary contact that led to the reinforcement of pre-mating barriers, the more recent clustering of coalescence rates on this region of scaffold 5 2000-3000 generations ago (Fig. 4e) may indicate when the reinforcement selection happened.

**Figure 4.**
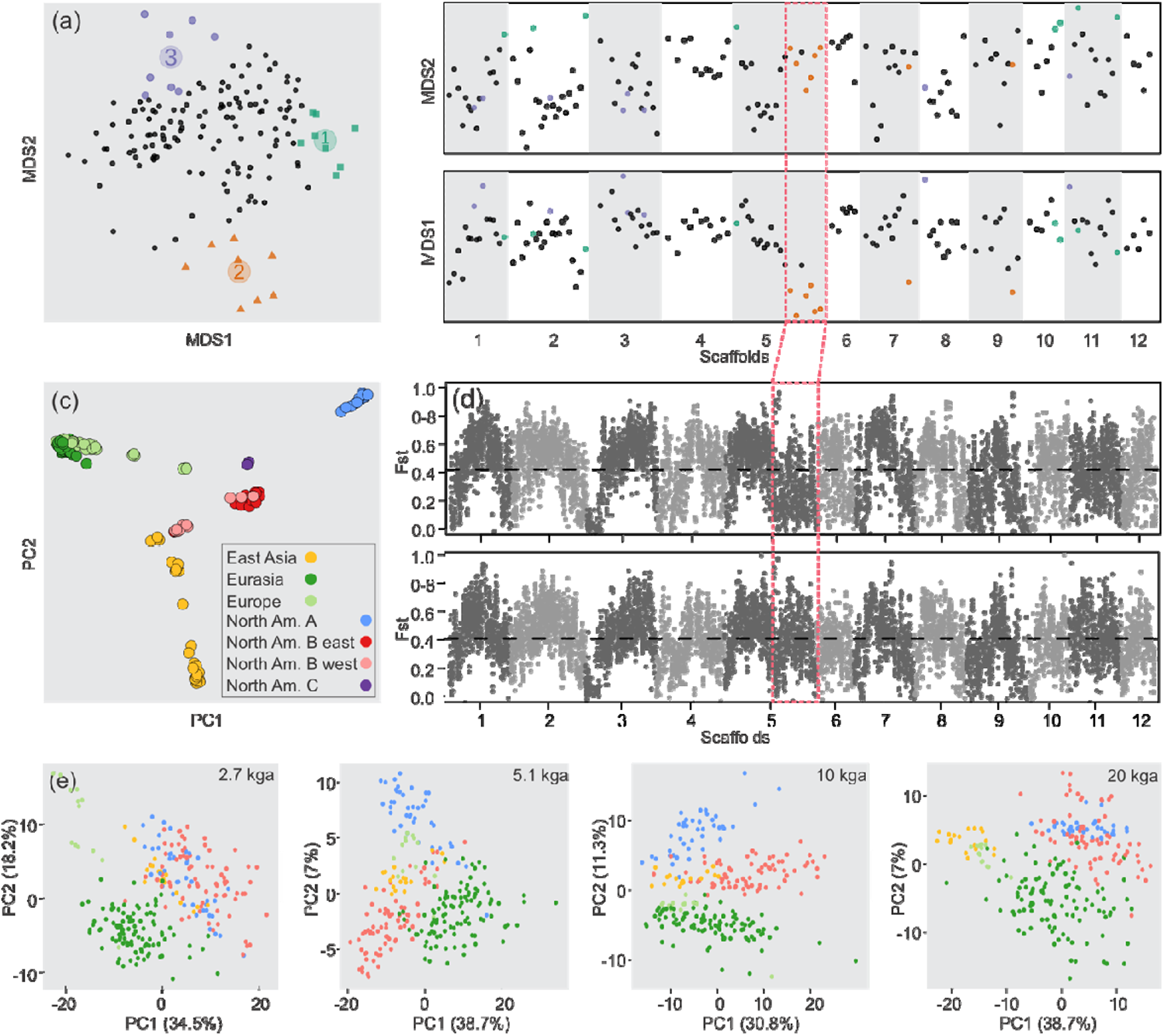
Scaffold 5 in the genome of Trichaptum abietinum show a distinct pattern **a**) Local PCA using Lostruct indicate three corners of genomic windows which deviate from the genomic average. **b**) The placement of the Lostruct outlier (corner) windows on MDS1 and MDS2 along the genome. **c**) A PCA of the SNPs in genomic windows of corner 2 show a different structure than the PCA of the genome average in Figure 1a). **d**) Fst genome scan of North America A and North America B compared to Fst of North America A and Eurasia. The second half of Scaffold5 which contains a high proportion genomic windows in corner 2 from the Lostruct analyses is boxed in. **e**) PCA of the coalescent rates of the second half of scaffold5 for selected time points.

### Concluding remarks

In this study we show how the morphospecies of a common forest fungus contain multiple differentiated and reproductively isolated lineages. These lineages have not evolved independently of each other, on the contrary, introgression and geneflow appear to be common. This reticulate pattern of evolution can be interpreted in light of the Pleistocene glacial cycles where previously isolated populations came into secondary contact during the interglacials. Several of the pre-mating barriers in *T. abietinum* fit a scenario of reinforcement, and we interpret the signals of secondary contact between reproductively isolated lineages and decrease of hybrid fitness with increased divergence, as additional support of this. Mating with the wrong lineage can be costly, nonetheless, we find large introgressed regions that have been retained, and we speculate that there could also have been certain adaptive advantages associated with inter-lineage matings. Future experiments on the ability of different hybrid dikaryons to produce fruit bodies with viable spores would allow an assessment of the relative strength of reproductive barriers at all stages of the life cycle and provide a more complete context to evaluate the case we make for reinforcement. Moreover, a controlled crossing scheme and sequencing of spore families from hybrid fruit bodies should be able to pinpoint the genomic basis of the premating barriers in *T. abietinum* and confirm the role of controlled cell death here.

Multiple reproductive barriers across different divergence levels make *T. abietinum* an exceptional study system to test outstanding questions in speciation genomics such as whether particular classes of genes or genetic elements have been involved repeatedly in independent speciation events. However, these qualities also make species concepts difficult to apply. It has been advised that biological species concepts in fungi are most useful in cases of sympatry^5^, whereas the phylogenetic species concept will be more discriminating in allopatry since significant phylogenetic divergence can precede the development of reproductive isolation in fungi^51,69,70^. In our case, North America A can be recognized as a distinct species based on both biological and phylogenetic species recognition criteria. Yet, North America A could, in theory, transfer their genetic material to intersterile lineages in North America and Asia through interfertile lineages in Europe. As such, *T. abietinum* resemble a ring species with North America A and B as intersterile end populations. Furthermore, following the criterion of reproductive isolation, the East Asian Yunnan sub-lineage should also be recognized as a distinct biological species. But consolidating a separate Yunnan species with the phylogenetic species concept means splitting up and recognizing both the other East Asian sublineages, and also North America B and Eurasia, as separate species in order to avoid paraphyletic species. It appears that fungal morphospecies, when analysed at a global scale, contain a multitude of lineages with nested levels of divergence and intersterility that are hard to fit within a rigid species concept.

## Materials and Methods

### Field sampling and culture isolation

To understand the global diversity and structure of the *T. abietinum* morphotaxon we collected fruit body samples in the field or had them sent in by collaborators between 2017 and 2020, where our aim was to cover as much of the known distribution range as possible. Individual collections were made from separate logs when possible, or in the cases where they were sampled from the same log, collections had to be at least 2 m apart to count as separate samples. Whenever possible, we included five samples from each locality. Monospore cultures were isolated by rewetting the dry fruit bodies in sterile H_2_O over night before they were attached to a petri dish lid with silicone grease from Merck Millipore (Merck KGaA, Darmstadt, Germany) over a 3% malt extract agar plate containing 25 mg/L streptomycin, 100 mg/L ampicillin, 10 mg/L tetracycline and 1 mg/L benomyl to inhibit the growth of contaminants. The lids with fruit bodies were set to shoot spores for 3 hours to 2 days, and subsequently replaced with sterile lids. These plates were set to germination at 19°C in the dark for 3-10 days, and germinated spores were excised and placed on new 3% malt extract agar plates with antibiotics and benomyl. After 2 weeks, slides of the mycelia were examined under the microscope to confirm the absence of clamp connections, indicating that they were monokaryotic cultures. To confirm that the isolated cultures were indeed *Trichaptum,* PCR of a small piece of culture tissue was performed using primers ITS1 and ITS4^71^ with Phire Plant Direct PCR Kit (Thermo Fisher Scientific, Waltham, MA, USA) following the manufacturer’s recommendations: 10 ul Phire Plant PCR Buffer, 4.8 ul H_2_0, 4 ul of 5 mM primer, 0.2 ul of Phire hot start II DNA polymerase, and 1 ul DNA containing 1 mg mycelium diluted in 20 ul dilution buffer. Amplicons were Sanger Sequenced at GATC Biotech/Eurofins Genomics (Eurofins Scientific, Luxembourg City, Luxembourg) using the ITS4 primer, after enzymatic clean up with Exo-SAP-IT PCR Product Cleanup Reagent (Thermo Fisher Scientific, Waltham, MA, USA) applying a 10x dilution of the Exo-SAP-IT but otherwise following the manufactureŕs instructions. Sanger sequences were blasted to the Unite^72^ and NCBI databases^73^ to confirm species identity, and aligned in Geneious Prime 2019.0.4 (https://www.geneious.com/) to an ITS2 alignment from Seierstad et al.^31^ to ascribe tentative lineage grouping. All work on the fungal cultures were carried out under a ESCO class II biological safety cabinet (ESCO lifesciences Gmbh, Friedberg, Germany) to avoid contamination.

### DNA extraction and Sequencing

To prepare DNA for Illumina sequencing, monokaryotic cultures were grown for 2-3 weeks on sterilised nitex nylon (Sefar AG, Heiden, Switzerland) placed on top of 3% malt extract agar plates. Mycelial tissue was scraped off from plates and homogenized in 2 ml Lysing Matrix E tubes (MP Biomedicals, Santa Ana, CA, USA) on a FastPrep-24 (MP Biomedicals, Santa Ana, CA, USA) for 2 x 20 s at 4.5 m/s^2^. DNA was extracted using the E.Z.N.A HP Fungal DNA kit (Omega Bio-Tek, Norcross, GA, USA) supplemented with 30 ul RNaseA (Qiagen, Hilden, Germany) prior to homogenization. The quantity and quality of DNA extractions were assessed with the Invitrogen Qubit fluorometer (Thermo Fisher Scientific, Waltham, MA, USA) using the Invitrogen dsDNA BR assay kit (Thermo Fisher Scientific, Waltham, MA, USA), NanoDrop Microvolume Spectrophotometer (Thermo Fisher Scientific, Waltham, MA, USA) and on a 1% agarose gel with GelRed (Biotium, Fremont, CA USA). Illumina sequencing was performed on either Hiseq 4000 or Novaseq at the Norwegian Sequencing Centre with library preparation as described in Peris et al.^32^. Briefly, 1 µg of genomic DNA was sheared to a target size of 300-400 bp with a Covaris E220 system (Covaris Inc., Woburn, MA, USA), cleaned on a small volume Mosquito liquid handler (TTP labtech) with a 1:1 ratio of Kapa Pure beads (Roche, Basel, Switzerland). Genomic libraries were generated using Kapa Hyper library prep kit (Roche, Basel, Switzerland) and Illumina UD 96 index kit (Illumina, San Diego, CA, USA) to add barcodes. Samples were pair-end sequenced at a read length of 2 x 150 bp.

### Read processing, SNP-calling and filtering

Bioinformatic analyses were performed on the SAGA supercomputer as part of the Sigma2 the National Infrastructure for High Performance Computing and Data Storage in Norway. Illumina reads were examined with FastQC 0.11.8 (http://www.bioinformatics.babraham.ac.uk/projects/fastqc/) and visualized with MultiQC^74^. Samples with less than 1 million reads were excluded from further analyses. Reads were trimmed with TrimGalore 0.6.5 (https://github.com/FelixKrueger/TrimGalore) at q30 and mapped with BWA 0.7.17 -mem^75^ to the PacBio assembly of the isolate TA-1010-6-M1 of *T. abietinum* from Peris et al.^32^, which is from the North America B lineage and consists of 12 nuclear scaffolds. Mapped reads were converted from sam to bam file format and sorted using SAMtools 1.9^76^ and duplicates were marked with the MarkDuplicates module in GATK 4.1.4.0. SNPs were called with GATK4’s pipeline of joint variant discovery^77^ by running Haplotypecaller in GVCF mode with ploidy = 1 followed by GenomicsDBImport and GenotypeGVCFs to produce the GVCF-file containing the joint variants in the data. The GVCF-file was quality filtered using BCFtools 1.10.2^76^ following an approach outlined in Barth et al.^78^ setting the thresholds to exclude sites that have: FS > 40, MQRankSum < −5.0 || > 5.0, ReadPosRankSum < −4, QD < 2, MQ > 40, SnpGap = 10, number of indels = 0, number of alleles = 2, INFO/DP > meanDP/2, missing data <20% and no monomorphic SNPs. The efficiency of filtering was visually controlled in R 4.2.2, and samples with > 50% of sites missing were removed from the VCF-file. A separate GATK4 joint variant discovery pipeline was run on the mapped reads in diploid mode to assess the VCF-file with VCFtools 0.1.16 –het^79^. In the output of VCFtools –het, 10 samples with F values < 0.8 were removed from the VCF-file as they likely represent dikaryons due to the high levels of heterozygosity. The resulting VCF-file contained 353 samples *T. abietinum* and 5 samples of the sister clade *T. fuscoviolaceum*, and was used as basis for further filtering for the different population genomic analyses. In several of the analyses, sites with missing data were removed from the VCF-file using the command bcftools view -g ^∧^miss in BCFtools. To obtain subsets of the data, we used vcf-subset in VCFtools.

### Analyses of Population structure

We used Principal Component Analysis (PCA), Admixture and a Maximum Likelihood (ML) phylogeny to assess the population structure in our SNP data. For the PCA and Admixture analyses the 5 *T. fuscoviolaceum* samples were removed from the VCF-file prior to filtering at minor allele frequency (MAF) > 0.05 and to only retain biallelic SNPs in BCFtools. The VCF-file was then LD pruned in PLINK 1.9^80^ with r2 threshold of 0.1 calculated within 50 bp windows with a step size of 10 bp to retain 46470 out of 776952 SNPs. The PCA was performed in Eigensoft 7.2.1^81^. Admixture^37^ was run with cross validation for 2-10 k clusters. The R package scatterpie (https://github.com/GuangchuangYu/scatterpie) was used to plot the admixture proportions onto a world map. For the maximum likelihood phylogeny, our SNP dataset including 5 samples of *T. fuscoviolaceum* was converted to the phylip format with vcf2phylip.py (https://github.com/edgardomortiz/vcf2phylip). IQtree^82^ was run on the phylip file with the options -T AUTO -m GTR+ASC -alrt 1000 -B 1000 using only biallelic sites. The ASC flag accounts for the lack of invariant sites in the SNP data by providing ascertainment bias correction. The phylogeny from IQtree was visualised in Figtree (http://tree.bio.ed.ac.uk/software/figtree/). We used the main lineages found in the PCA, Admixture and ML phylogeny to ascribe samples to lineages for a Fst genome scan analysis. Here, a VCF-file without *T. fuscoviolaceum* and filtered in BCFtools to contain only biallelic SNPs was converted to a geno file with parseVCF.py. Sliding window genome wide statistics of Fst were calculated from the geno file popgenWindows.py using a window size of 10 kb and step size of 5 kb. Both scripts are available at https://github.com/simonhmartin/genomics_general. The admixed samples in central Europe and the North America C were not included in the analysis due to high admixture proportions and low sample size, respectively. A separate Fst genome scan was run at finer scale where the North America A, North America B, and East Asia was also split up into sublineages. Tajimás D and pi were calculated in the R package PopGenome^83^.

### Analyses of Recombination

Coalescence analysis in Relate requires estimates of the recombination rate, and we used LDhelmet^84^ to obtain estimates of the recombination rate along each of the 12 nuclear scaffolds. We performed this analysis on 53 North America B samples belonging to the same lineage and population (New Brunswick) as the reference strain TA-1010-6-M1. A VCF-file containing the 53 samples filtered to include only biallelic SNPs and with all missing sites removed with BCFtools was split into individual scaffolds with vcftools --chr. Each VCF-file was then converted to the fasta input file for LDhelmet with python scripts. LDhelmet was run as recommended in the manual with the steps: find_confs -w 50, ldhelmet pade -t 0.01 -x 11, ldhelmet rjmcmc -p output.pade -w 50 -b 50.0 and using a linkage table generated with LDhat ^85^, and finally extracting the mean output with ldhelmet post_to_text. The linkage table was generated in LDhat complete with the parameters -n 53 -rhomax 100 -n_pts 101-theta 0.01, and converted to the LDhelmet format using ldhelmet convert_table --input_table. We obtained our estimate of theta with the R package Pegas^86^ using scaffold 12 of the same VCF-file of the 53 North America B samples. Scaffold 12 is the smallest scaffold and was chosen to make the computation manageable.

### Coalescence analysis

Coalescence analyses were performed with Relate^39^ to investigate population histories of the major lineages. The VCF-file without *T. fuscoviolaceum* and filtered to contain only biallelic SNPs and no missing sites in BCFtools were split up into VCF-files for each individual scaffold with VCFtools --chr. The input haps files for Relate were generated from these VCF files using PLINK/2.00 --export haps^80^. The resulting haps files were then converted to reflect the haploid nature of our sequence data by excising only every other column for each sample. RelateFileFormats -mode FlipHapsUsingAncestor was used to infer ancestral allele state using a consensus sequence of *T. fuscoviolaceum.* The consensus sequence was generated with vcf-consensus in VCFtools using the PacBio reference assembly of *T. abietinum* and extracting the 5 samples of *T*. *fuscoviolaceum* from the general VCF-file. Relate was then run in --mode All setting the mutation rate to 1e-7 as studies on another rapidly fruiting wood decay fungus found a mutation rate within this order of magnitude^87^. The effective population size (Ne) was set to 10,000, and the recombination map for the Relate analysis was obtained by multiplying the population recombination rate from LDhelmet with 100/(4*Ne), The resulting anc and mut files of each scaffold from the Relate analysis were used to re-estimate coalescence rates and effective population sizes with the Relate script EstimatePopulationSize.sh. Here we combined all scaffolds and specified -- num-iter 1 to avoid bias, --threshold 0 to include all sites, and --years_per_gen 1 to specify a generation time of 1 year. EstimatePopulationSize.sh was run separately to obtain the average within coalescence rate of each population that was plotted Fig.1e and used to estimate changes in effective population size for each generation, and also run with all samples without any population specification to obtain coalescence rates between all samples. Principal components of these coalescence rates between all samples were computed with prcomp() in R with normalization of the columns prior to plotting in R. A test for positive selection was calculated on the inferred genealogy for each major lineage separately based on the DetectSelection.sh in Relate on the re-estimated anc and mut files. The North America C lineage was excluded from the analysis due to low sample size (n=2).

### Mating experiments

In order to test if monokaryotic isolates could mate and form dikaryotic mycelia, paired isolates were placed 4 cm apart on 3% malt extract agar plates. When the two mycelia had grown out to encounter each other, after 2-3 weeks, a small piece from the contact zone was excised and re-plated on a new 3% malt extract agar plate. Microscope slides were made of this new culture after one week of growth, and examined under a Nikon Eclipse 50i microscope (Nikon Instruments Europe BV, Amsterdam Netherlands) at 400x magnification to assess the frequency of clamps. Each slide was scored for the proportion of clamps present on the first 10 observed septae (cell junctions). If none of the first 10 septae contained clamps, different areas of the slide were also searched, and if clamps were then detected a note of this was made. The majority of crosses were set up in triplicates, and mating success was evaluated as average proportion septa with clamps across replicates. In the classification of matings used for figure 2, matings were rated as positive if on average 5/10 or more of the septae contained clamps, intermediate if 2/10-4/10 contained clamps, marginal if 1/10 or less contained clamps and in cases were none of the first 10 examined septae contained clamps, but clamps were found in other parts of the slide, and negative if no clamps were found at all. Images of the slides were acquired with a Zeiss Axioplan-2 imaging with Axiocam HRc microscope camera (Zeiss, Oberkochen Germany). The ability to mate between groups were correlated to their average genome wide Fst distance with a Pearsońs correlation test.

### Dikaryon fitness experiment

As an assessment of the fitness of dikaryons, 2 cm x 2 cm x 0.5 cm woodblocks of *Pinus sylvestris* placed on top of petri dishes with water agar and inoculated with dikaryons. Decay was measured after 3 months at 19 **°**C by calculating percentage mass loss in the woodblock. For the experiment we used four hybrid dikaryons between the North America A, North America B, Eurasian, European, and East Asian lineages, and eight dikaryons from within each of these lineages. For the within lineage dikaryons, both parents were from the same or nearby population. Each dikaryon was set up in four replicates. The average difference of mass loss between the parental lineages and the hybrid in each case were correlated to Fst distances with a Pearsońs correlation test. T-tests in R were used to determine if the wood decay ability of the hybrid dikaryons were significantly different than dikaryons of each parent lineage.

### Genome Wide Association Studies

We carried out a genome wide association study (GWAS) to associate intersterility with North America A to our SNP data. The phenotype in our GWAS was set as binary according to intersterility with the North America A lineage. For crosses where no clamps or only a few clamps were formed (≤ 1/10) phenotype was set as 0, and for the rest phenotype was set as 1. Intersterility for the samples where we did not have crossing information were inferred based on their lineage, so that it was 0 for North America B, North America C, East Asia SW and East Asia SC. Phenotype was set as 1 for the samples belonging to the Eurasia, Europe, East Asia NE and East Asia N lineages. Map and raw files were produced using PLINK2.00 -- recode on VCF file without *T. fuscoviolaceum* filtered for missing sites in BCFtools. The GWAS was run with the amm_gwas script in R available at https://github.com/arthurkorte/GWAS/blob/master/gwas.r. To account for population structure a VanRaden.2 kinship matrix was calculated with the gMatrix package available at https://github.com/filippob/introduction_to_gwas/blob/master/software/gMatrix_0.2.tar.gz. To leverage our crossing data and account for intersterility with more than 1 lineage, a subset of the 25 samples for which we had exact crossing information was given to SCOPA^57^ for a multivariate analysis to detect correlated phenotypes. A gen file was generated with PLINK2.00 from a VCF-file of the 25 samples with the --export oxford setting as input for the SCOPA analysis. As phenotype, we used intersterility with 12 of the 25 samples to reduce the computational time. A given sample were set to be able to mate with itself for the analysis, as this is governed by the homogenic incompatibility system of the mat-A and mat-B loci that control mating within an ISG, whereas we were interested in detecting the genes that cause intersterility between ISGs.

### ΔFst

ΔFst was calculated for 10 kb Fst genome scan windows of intersterile lineages as contrasted to interfertile lineages. A high positive ΔFst thus indicates a region that is highly differentiated between lineages that cannot mate, but similar in the lineages that can mate. We selected our comparisons where the interfertile and intersterile comparisons would involve the same lineage, and where the intersterile and interfertile lineage compared to it would be as closely related to each other as possible. We identified the 95% percentile outlier windows towards the positive tail end of the empirical distribution of all ΔFst windows for each of the 17 comparisons.

### Gene function

We used the annotation of our reference genome from^32^ as basis of gene function searches. Metacaspases and the pfam domains PF05729 and PF00400, which are associated with het-genes in the sister phylum Ascomycota, were pulled out from the gff file containing the annotations. Tblastx searches for het gene homologs were performed using het gene sequences in *Podospora anserina* and *Neurospora crassa* downloaded from the uniprot database^88^ as described in^61^. An enrichment analysis of GO terms within the outlier ΔFst windows, GWAS snps, and SCOPA snps, was conducted in R with Rtracklayer available at https://bioconductor.org/packages/release/bioc/html/rtracklayer.html and topGO available at https://bioconductor.org/packages/release/bioc/html/topGO.html following the setup on: https://ucdavis-bioinformatics-training.github.io/2018-June-RNA-Seq-Workshop/friday/enrichment.html. For the p-value, a Benjamini-Hochberg corrected threshold of p = 0.05 was used. Similarly, enrichment analyses of PFAM domains falling within genes with significant GWAS SNPs, SCOPA SNPs, and within ΔFst windows were carried out in R. In the enrichment analyses, we considered only the subset of pfams that occurred in our region of interest. The test was set up as described in^89^ with Fisher’s exact test, and Benjamini-Hochberg corrected threshold of p = 0.05.

### Treemix analysis

Treemix infers a maximum likelihood tree including an optional number of m migration edges between branches and thereby can account for hybridization and introgression between lineages. The linkage pruned VCF file from our PCA and Admixture analysis was converted to a strat file using plink2treemix.py (https://github.com/thomnelson/tools/blob/master/plink2treemix.py) and vcf2tremix (https://github.com/speciationgenomics/scripts) for Treemix^63^. Treemix analysis was run with -m edges of 0-10 and a block size of -k 500. Results were evaluated with the R package OptM ^90^.

### Demographic modelling

For the demographic modelling, a subsampled VCF-file filtered for missing sites of 13 samples each from the North America A, North America B, European and Yunnan population of the East Asian lineages covering a similar geographic range were chosen. The model of coalescence order between the lineages we used in these analyses were guided by the ML phylogeny, coalescence analysis, and Treemix, which all indicate North America A as the first lineage to diverge. We included only the most realistic scenarios of geneflow, namely across the Bering strait between North American groups and East Asia, between Eurasia and East Asia, and within the North American groups. Gene flow between North America and Europe, i.e. across the Atlantic, was not considered, with the exception of a simple model with only one event of gene flow. Site frequency spectrum files were generated from the VCF-file with easySFS -a -f (https://github.com/isaacovercast/easySFS). For each demographic model, Fastsimcoal 2.6^65^ was initiated with 50 independent runs with -m -n 100000 -L 50 -s 0 -M, which specifies 50 cycles of 100,000 coalescent simulations per cycle. The mutation rate in the tpl files was set to 1e-7, the same as in our coalescence analysis in Relate. The run with the highest likelihood and its estimated parameters for each model was extracted with scripts available on https://github.com/speciationgenomics/scripts. A likelihood distribution for each model was obtained by running fsc26 -n 1000000 -m -q with the best parameter values for the given model with 100 iterations and collecting the likelihood values for each run using a script available at https://github.com/speciationgenomics/scripts. The likelihood values were then used to compute AIC-distributions of the models, and both were plotted in R. An initial comparison of models with 1, 2 or 3 splits between the lineages identified a model with 2 splits as the model with best support. This model was used as basis for further testing of models with gene flow.

### Local PCA and Coalesence Analysis of Scaffold 5

Our local PCA was run with Lostruct^67^ using scripts available at https://github.com/petrelharp/local_pca/blob/master/templated/README.md. The analysis was run for 5 kb windows on a 0.05 MAF-filtered and biallelic VCF-file. The second half of scaffold 5 was extracted using VCFtools --from-bp 3213128 --to-bp 6426255, and analysed separately in Relate in the same manner as the other analyses in Relate described under the section on coalescence analysis.

### Data visualization

If not otherwise mentioned, data was visualized in R with ggplot2^91^, and final graphic design adjustments were made in CorelDRAW (Corel corporation, Ottawa, Canada).

## Supporting information

Table1, Supplementary figure dikaryon fitness, Supplementary figure GWAS, Supplementary figure Fastsimcoal2, Supplementary table2,Supplementary table3

## Acknowledgements

This work was supported by the Research Council of Norway (RCN) grant No. RCN 274337 and the University of Oslo and UiO:Life Science internationalization support. David Peris is supported by the RCN grant No. RCN 324253, and the Generalitat Valenciana plan GenT grant No. CIDEGENT/2021/039. We thank all the sample collectors who made this study possible. The computations were performed on resources provided by UNINETT Sigma2 – the National Infrastructure for High Performance Computing and Data Storage in Norway.

