## Supplementary material for "Reticulate evolution and rapid development of reproductive barriers upon secondary contact pose challenges for species delineation in a forest fungus": Table1, Supplementary figure dikaryon fitness, Supplementary figure GWAS, Supplementary figure Fastsimcoal2, Supplementary table2,Supplementary table3

**Table1.** Parameter estimates and likelihood values of different demographic models in fastsimcoal2. All time estimaes such as T.coal and T.mig are given in generations. The best likelihood and AIC values are indicated with *.

| **Model name** | **Mig1** | **Mig2** | **Mig3** | **Mig4** | **T.coal1** | **T.coal2** | **T.mig1** | **T.mig2** | **T.mig3** | **T.mig4** | **Best likelihood** | **AIC** |
| --- | --- | --- | --- | --- | --- | --- | --- | --- | --- | --- | --- | --- |
| nomig |  |  |  |  | 104488 | 81119 |  |  |  |  | -4961578 | 22848932 |
| mig1 | NoramA  Euro |  |  |  | 87143 | 57668 | 1351 |  |  |  | -4955439 | 22820669 |
| mig2 | NoramA NoramB |  |  |  | 12311 | 6862 | 397 |  |  |  | -4950017 | 22795698 |
| mig3 | NoramA EastAsia |  |  |  | 37300 | 23243 | 787 |  |  |  | -4956527 | 22825677 |
| mig4 | NoramA NoramB | NoramB EastAsia |  |  | 100135 | 51084 | 1458 | 1718 |  |  | -4928962 | 22698743 |
| mig5 | NoramA NoramB | NoramA EastAsia |  |  | 74668 | 22727 | 1741 | 372 |  |  | -4935361 | 22728210 |
| mig6 | NoramA NoramB | Euro EastAsia |  |  | 59751 | 38789 | 1117 | 495 |  |  | -4936567 | 22733766 |
| mig8 | NoramA EastAsia | NoramB EastAsia |  |  | 71474 | 45082 | 720 | 1087 |  |  | -4929747 | 22702356 |
| mig9 | NoramA EastAsia | Euro EastAsia |  |  | 44626 | 25785 | 386 | 814 |  |  | -4941326 | 22755681 |
| mig10 | NoramA EastAsia | NoramB Euro |  |  | 100374 | 64441 | 769 | 4743 |  |  | -4940129 | 22750170 |
| mig7 | NoramA NoramB | NoramB EastAsia | NoramA EastAsia |  | 94768 | 63572 | 812 | 955 | 58716 |  | -4930303 | 22704922 |
| mig11 | NoramA NoramB | NoramB EastAsia | NoramB Euro |  | 51552 | 27793 | 1430 | 1012 | 21812 |  | -4932762 | 22716247 |
| mig13 | NoramA EastAsia | NoramB EastAsia | Euro EastAsia |  | 50295 | 30308 | 353 | 34991 | 1005 |  | -4941292 | 22755527 |
| mig12 | NoramA NoramB | NoramB EastAsia | NoramA EastAsia | Euro  EastAsia | 113274 | 59875 | 4864 | 3467 | 2082 | 1104 | -4902064* | 22574887* |

**Supplementary figure dikaryon fitness**

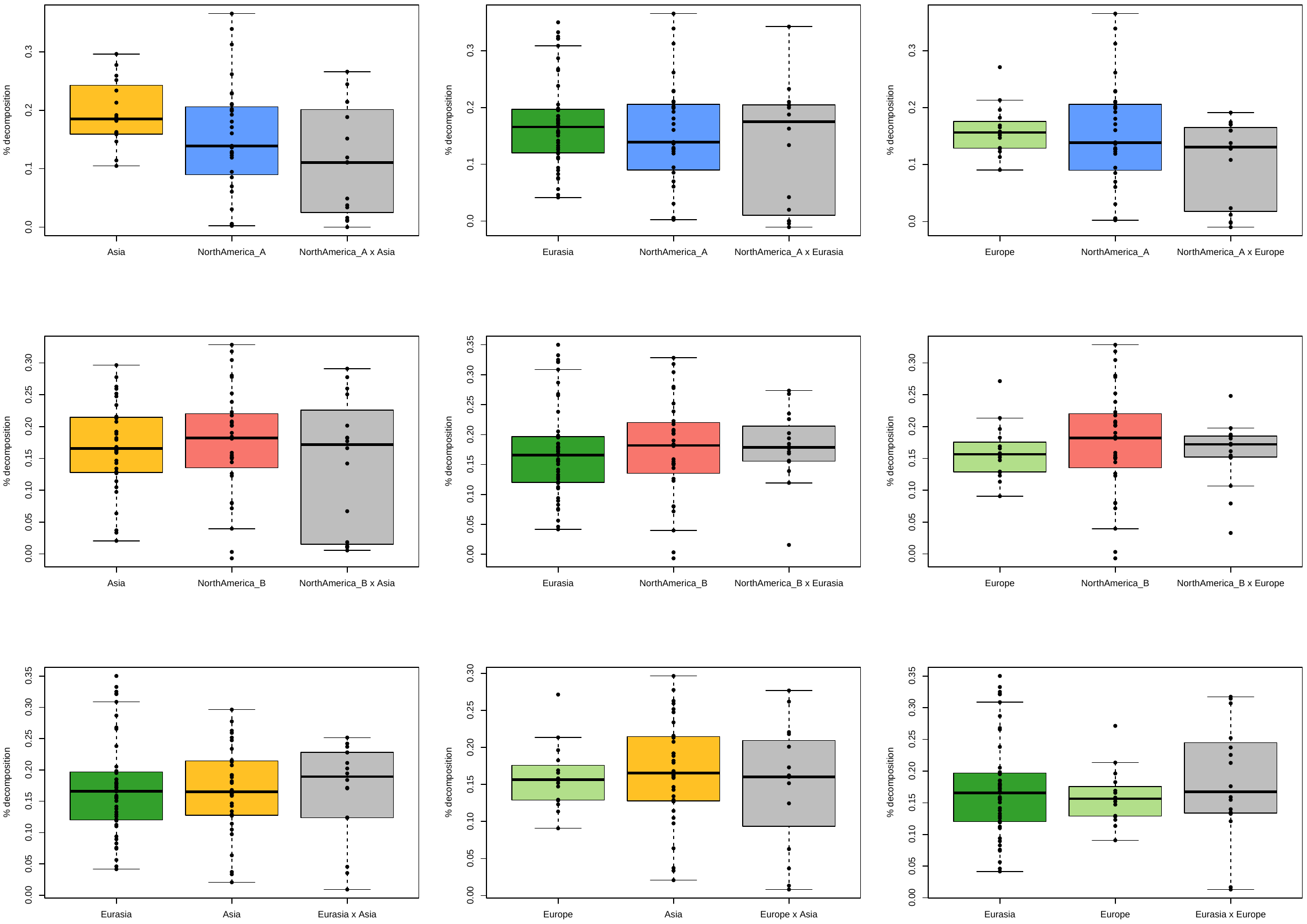

**Supplementary figure GWAS**

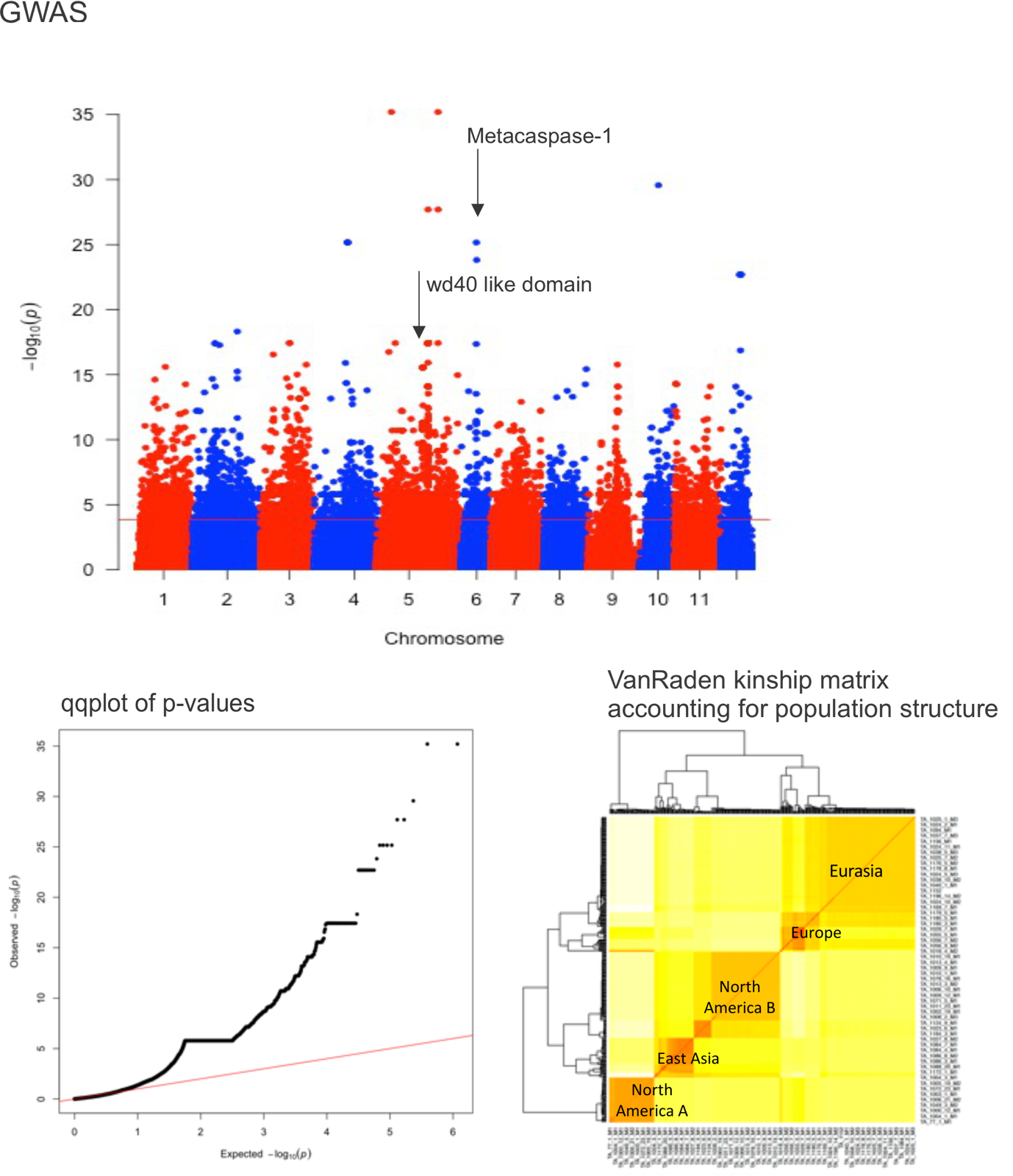

**Supplementary figure Fastsimcoal2**

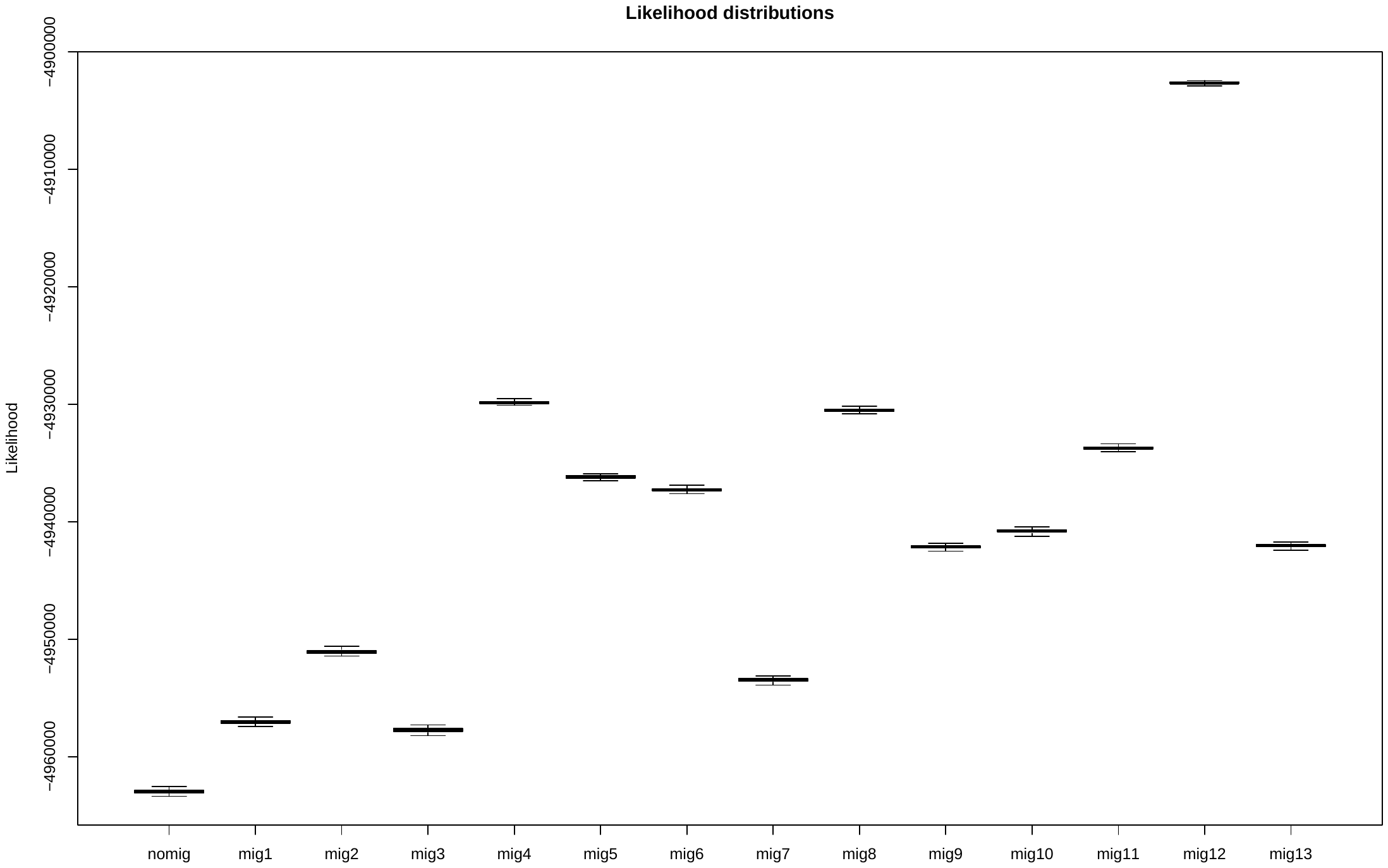

**Supplementary table2**. Genome wide average of Fst between sublineages.

|  | North Am. A E | North Am. A S | North Am. B E | North Am.B W | East Asia SW | East Asia SC | East Asia NE | East Asia N | Eurasia |
| --- | --- | --- | --- | --- | --- | --- | --- | --- | --- |
| **North America A S** | 0.07 |  |  |  |  |  |  |  |  |
| **North America B E** | 0.42 | 0.21 |  |  |  |  |  |  |  |
| **North America B W** | 0.46 | 0. 44 | 0.13 |  |  |  |  |  |  |
| **East Asia SW** | 0.41 | 0.44 | 0.21 | 0.35 |  |  |  |  |  |
| **East Asia SC** | 0.29 | 0.42 | 0.10 | 0.24 | 0.14 |  |  |  |  |
| **East Asia NE** | 0.49 | 0.45 | 0.28 | 0.41 | 0.10 | 0.18 |  |  |  |
| **East Asia N** | 0.31 | 0.43 | 0.08 | 0.20 | 0.16 | 0.11 | 0.16 |  |  |
| **Eurasia** | 0.41 | 0.17 | 0.26 | 0.19 | 0.18 | 0.08 | 0.25 | 0.07 |  |
| **Europe** | 0.41 | 0.43 | 0.29 | 0.45 | 0.41 | 0.34 | 0.45 | 0.35 | 0.17 |

|  | **North Am. A** | **North Am. A S** | **North Am. B east** | | **North Am. B west** | **East Asia N** | **East Asia**  **NE** | **East Asia**  **SC** | **East Asia**  **SW** | **Eurasia** | **Europe** |
| --- | --- | --- | --- | --- | --- | --- | --- | --- | --- | --- | --- |
| **π** | 0.0014 | 0.0015 | | 0.0015 | 0.0011 | 0.0129 | 0.0088 | 0.0118 | 0.0017 | 0.0016 | 0.0014 |
| **Tajima´s D** | 0.279 | -0.420 | | 0.090 | -0.202 | 0.091 | 0.180 | -0.458 | -0.029 | -0.153 | 0.239 |

**Supplementary table3.** Nucleotide diversity **π** and Tajima´s D estimated in PopGenome from the SNP data.
